# Host-derived protease promotes aggregation of *Staphylococcus aureus* by cleaving the surface protein SasG

**DOI:** 10.1101/2022.12.13.520364

**Authors:** Heidi A. Crosby, Klara Keim, Jakub M. Kwiecinski, Christophe J. Langouët-Astrié, Kaori Oshima, Wells B. LaRivière, Eric P. Schmidt, Alexander R. Horswill

**Author notes:** **Corresponding author:** Alexander R. Horswill, Ph.D., University of Colorado School of Medicine, Department of Immunology and Microbiology, 12800 E. 19th Ave., RC1N-9101, Mail Stop 8333, Aurora, CO 80045. These authors contributed equally (Heidi A. Crosby, Klara Keim).

## Abstract

*Staphylococcus aureus* is one of the leading causes of hospital acquired infections, many of which begin following attachment and accumulation on indwelling medical devices or diseased tissue. These infections are often linked to establishment of biofilms, but another often overlooked key characteristic allowing *S. aureus* to establish persistent infection is formation of planktonic aggregates. Such aggregates are physiologically similar to biofilms and protect pathogen from innate immune clearance and increase its antibiotic tolerance. The cell wall-associated protein SasG has been implicated in biofilm formation via mechanisms of intercellular aggregation, but the mechanism in the context of disease is largely unknown. We have previously shown that expression of cell wall-anchored proteins involved in biofilm formation is controlled by the ArlRS-MgrA regulatory cascade. In this work, we demonstrate that the ArlRS two-component system controls aggregation, by repressing expression of *sasG* by activation of the global regulator MgrA. We also demonstrate that SasG must be proteolytically processed by a non-native protease to induce aggregation, and that strains expressing functional full-length *sasG* aggregate significantly upon proteolysis by a mucosal-derived host protease found in human saliva. We used fractionation and N-terminal sequencing to demonstrate that human trypsin within saliva cleaves within the A domain of SasG to expose the B domain and induce aggregation. Finally, we demonstrated that SasG is involved in virulence during mouse lung infection. Together, our data point to SasG, its processing by host proteases, and SasG-driven aggregation as important elements of *S. aureus* adaptation to host environment.

## Introduction

*Staphylococcus aureus* asymptomatically colonizes the nostrils, throat, and skin of ∼30% of the population, and a portion also carry *S. aureus* in their oral cavity [1-5]. Nasal carriage is a significant risk factor for developing nosocomial infections [6, 7], with ∼80% of infections caused by the patient’s colonizing strain [8-10]. *S. aureus* is one of the leading causes of healthcare-associated infections, such as surgical site infections and central line-associated bloodstream infections [11], imposing a substantial burden on the healthcare system. While these infections are often challenging to treat, the rise of methicillin-resistant *S. aureus* (MRSA), which causes over 119,000 of these infections annually in the US, has further exacerbated treatment challenges and increases healthcare costs by nearly one billion dollars annually [12-15].

*S. aureus* is one of the most prevalent pathogens in chronic wound infections [16-18], and is one of the first pathogens to colonize in the cystic fibrosis (CF) lung [19]. The occurrence of chronic and persistent *S. aureus* infections is in part due to aggregation mechanisms and the ability of this pathogen to adhere to indwelling medical devices as a biofilm [20, 21]. However, in the absence of an implanted medical device, *S. aureus* can form free-floating aggregates that are physiologically similar to biofilms and are likewise more antibiotic resistant [22, 23]. It has been suggested that bacterial aggregates predominate in chronic infections such as osteomyelitis [24], chronic wounds [25], and in the lungs of cystic fibrosis (CF) patients [26, 27]. Intensive efforts to clear MRSA lung infections in CF patients, sometimes using up to five different antibiotics, has shown some promise, although ∼15% of patients still harbor MRSA at the end of the intervention period [28-30]. A better understanding of *S. aureus* biofilm formation and aggregation may lead to alternative therapies for these difficult to treat infections.

MRSA aggregation observed in clinical infections has been described as groups of closely attached cells that are not surface attached, and similarly to mature biofilms they provide protection from environmental stress and allow for persistence [22].

Aggregates and biofilms are difficult to treat in part because they are up to 1000-fold more resistant to antibiotics than planktonic cells [22, 31, 32]. This increased tolerance is thought to be due to a combination of slowed diffusion of antibiotics through the extracellular matrix and slower growth of cells within the community of cells [33]. In addition, aggregates are more resistant to clearance by the innate immune system, in part due to their large size, which impedes phagocytosis, and their ability to secrete and concentrate toxins that target leukocytes [34-37].

One of the key drivers of biofilm formation and aggregation in *S. aureus* is the large, cell wall-attached surface protein G (SasG) [38-40]. SasG, and its *S. epidermidis* homolog Aap, consist of multiple domains with distinct functions (**Fig. 1A**). The A domain, which has 59% identity to Aap, is implicated in binding to corneocytes [41] and nasal epithelial cells [42], and has a short, variable repeat region and an L-type lectin subdomain. In full-length SasG, the B domain, which has 60-67% identity to Aap depending on B-repeat number, consists of 2-17 repeats of alternating G5 subdomains and E spacers [38, 43, 44]. These G5-E repeats can dimerize in a Zn-dependent manner to form a twisted cable structure that facilitates intercellular interactions [45]. In *S. epidermidis*, the Aap A domain is removed by the metalloprotease SepA, allowing the exposed B domains to dimerize and promote biofilm accumulation [46]. Exogenous addition of the host proteases trypsin and cathepsin G can also enhance *S. epidermidis* biofilm formation through processing of Aap [43]. Whether SasG also needs to be proteolytically processed is not known, although it appears that none of the known proteases secreted by *S. aureus* can specifically target SasG [38].

**Fig. 1.**
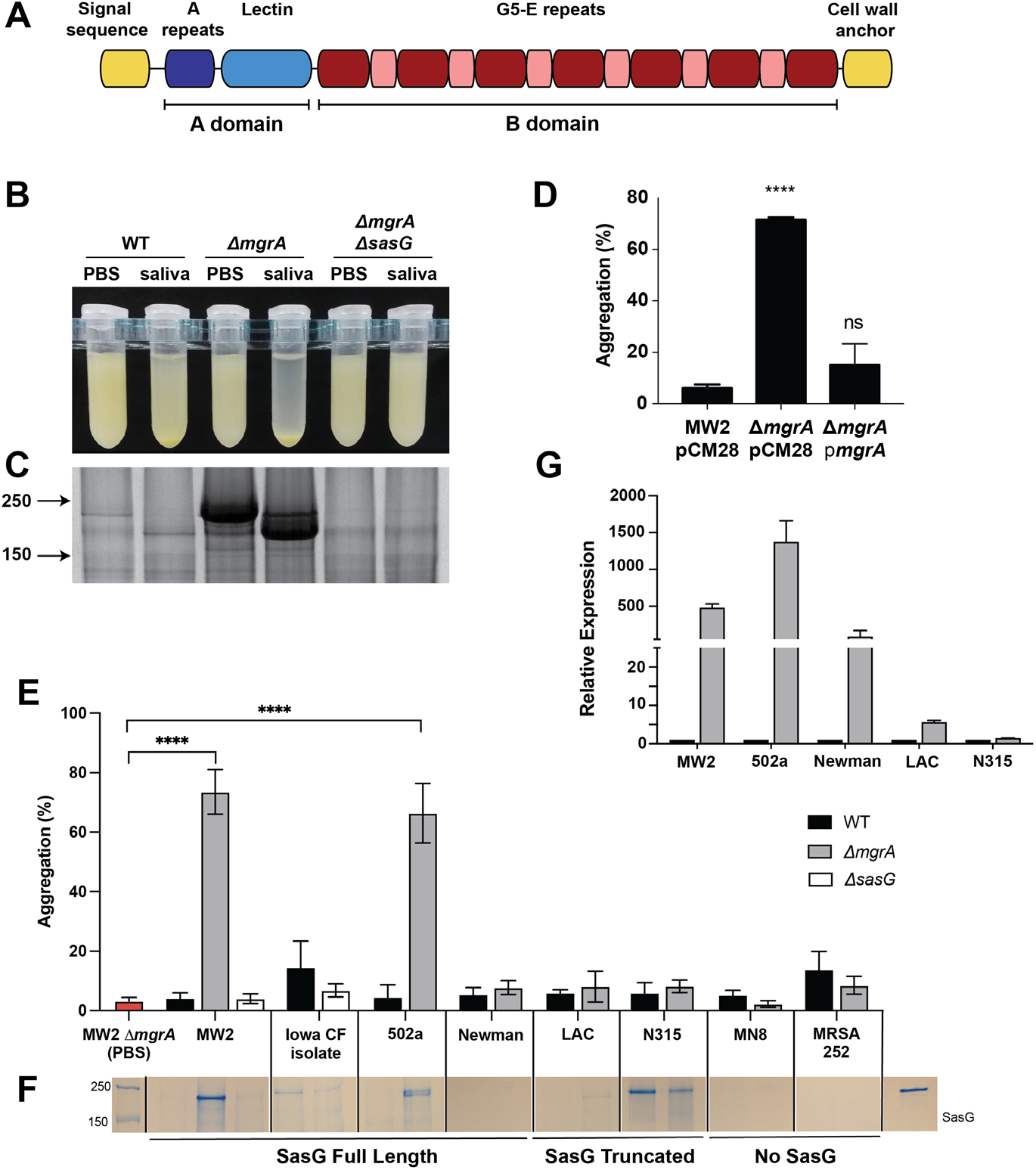
*S. aureus* aggregates in presence of human saliva and high SasG levels. (A) Schematic of SasG domains. (B-D) Overnight cultures of the indicated MRSA MW2 strains were spun down and resuspended in either phosphate buffered saline or clarified human saliva. (B) Photo shows aggregation of the Δ*mgrA* mutant after one hour of incubation at room temperature. (C) Coomassie stained SDS-PAGE gel shows cell wall preps from these same samples after one-hour incubation as described above. Experiment is representative of at least three replicates. (D) Quantification of aggregation of MW2 WT with the empty vector pCM28, or Δ*mgrA* mutant with either pCM28 or the complementation vector pCM28-*mgrA* (pHC66) in the presence of saliva. Data represent averages and standard deviations of three separate experiments. Statistical significance was calculated by One-way ANOVA. ****, p ≤ 0.0001; ns, not significant. (E) Various *S. aureus* strains with full-length, truncated or lacking *sasG* were incubated with human saliva and aggregation was measured following 2hrs of incubation. (F) Cell wall proteins were precipitated from overnight cultures and run on SDS PAGE to observe relative SasG expression levels. (G) Quantification of *sasG* gene expression of various *S. aureus mgrA* mutant strains relative to the respective wild-type *sasG* expression (n=3). Values are normalized to *gyrB* expression in each strain.

Expression of *sasG* is variable across *S. aureus* clinical isolates. SasG is constitutively expressed by some clinical isolates [47], and the presence of anti-SasG human antibodies demonstrates its expression during infection [48, 49]. However, commonly used laboratory strains either lack functional SasG, or do not express it under laboratory conditions [47, 50]. Recently, it has become apparent that this lack of SasG expression might be due to its repression by an ArlRS – MgrA regulatory cascade under *in vitro* conditions [49, 51].

In this project, we took advantage of the high level of SasG expression in a *S. aureus* Δ*mgrA* strain to investigate the role of SasG in aggregation and virulence. We identified that the presence of SasG increases *S. aureus* virulence during lung infection, and that the cleavage of the N-terminal portion of the A domain of SasG is necessary for *S. aureus* to aggregate. Since *S. aureus* does not appear to cleave SasG on its own, SasG cleavage during infection must be mediated by host proteases. Such cleavage leads to SasG-mediated aggregation of *S. aureus*, which is reflected as increased virulence of SasG-expressing strain during lung infection. Overall, the host-driven cleavage of SasG establishes an unusual and novel way of sensing and responding to the host environment.

## Results

### SasG saliva interaction and expression levels across S. aureus strains

Aspiration of saliva is often a precursor to lung infections [52-55], leading us to investigate how MRSA reacts to the presence of human saliva. We made a somewhat surprising observation that a USA400 MRSA Δ*mgrA* mutant strain aggregated to high levels when the cells were resuspended in human saliva, while the WT strain remained in suspension (**Fig. 1B**). Knowing there is differential surface protein expression in Δ*mgrA* mutants [49], we ran Coomassie protein gels (**Fig. 1C**) and observed dramatic salivary processing of a large protein that we reasoned might be surface protein G (SasG). Upon constructing a MRSA Δ*mgrA* Δ*sasG* double-mutant, the protein and aggregation phenotype both disappeared (**Fig. 1B,C**), demonstrating this phenotype is due to SasG. Additionally, the aggregation could be complemented by providing *mgrA* on a plasmid (**Fig. 1D**).

We next investigated the generality of this phenotype in *S. aureus*. We compared sequenced *S. aureus* strains containing functional chromosomal copies of *sasG* including community-acquired MRSA (CA-MRSA) USA400 strain MW2, Newman, 502a, and a CF clinical MSSA isolate AH4654. We also included strains that expressed a truncated form of SasG such as those of USA300 strain LAC and USA100 strain N315. Finally we included strains lacking a copy of the *sasG* gene altogether, such as USA200 strains MN8 and MRSA252, as controls for comparison. Strains with a functional, full-length version of SasG protein exhibited high levels of saliva-induced aggregation in the absence of *mgrA* (**Fig. 1E**) and we observed abundant SasG in cell wall preparations (**Fig. 1F**). The CF clinical isolate AH4654 exhibited lower expression levels and intermediate aggregation (**Fig. 1E**), although the genetic composition is almost identical to MW2, the functionality of the ArlRS-MgrA system in relation to SasG is not clear in this strain. Unexpectedly, Newman exhibited no visible expression of SasG protein (in WT or Δ*mgrA* mutant) and little aggregation despite having a full-length version of SasG encoded in the genome (**Fig 1E, F**). N315 expressed a protein of size to SasG but did not clump at all. These data were confirmed by qPCR quantifying *sasG* expression **(Fig. 1G**). In general, our observations indicate that *S. aureus* strains with a full-length SasG, under conditions that induce *sasG* gene expression, will aggregate in the presence of human saliva.

### Molecular details of MgrA repression of sasG gene

To investigate transcriptional control of *sasG* in the (CA-MRSA) USA400 strain MW2, we constructed a P_*sasG*_-sGFP reporter plasmid (pHC127) with *sasG* promoter fused to a gene encoding sGFP. This plasmid was transformed into mutants of the ArlRS and MgrA regulatory systems, previously suspected to repress the expression of SasG, and the expression levels were monitored over 24 h (**Fig. 2A**). The highest expression was observed in the Δ*mgrA* mutant, followed by the Δ*arlRS* mutant, with minimal expression in WT. The high expression in Δ*mgrA* mutant was confirmed at the protein level (**Fig. 2B**). We analyzed the *sasG* promoter region by 5’RACE to identify a putative housekeeping promoter and transcriptional start site (**Fig. 2C**). Putative MgrA repressor binding sites are shown that overlap the promoter region. Overall, our findings confirms that expression of SasG in laboratory growth media is repressed by the ArlRS – MgrA regulatory cascade.

**Fig. 2.**
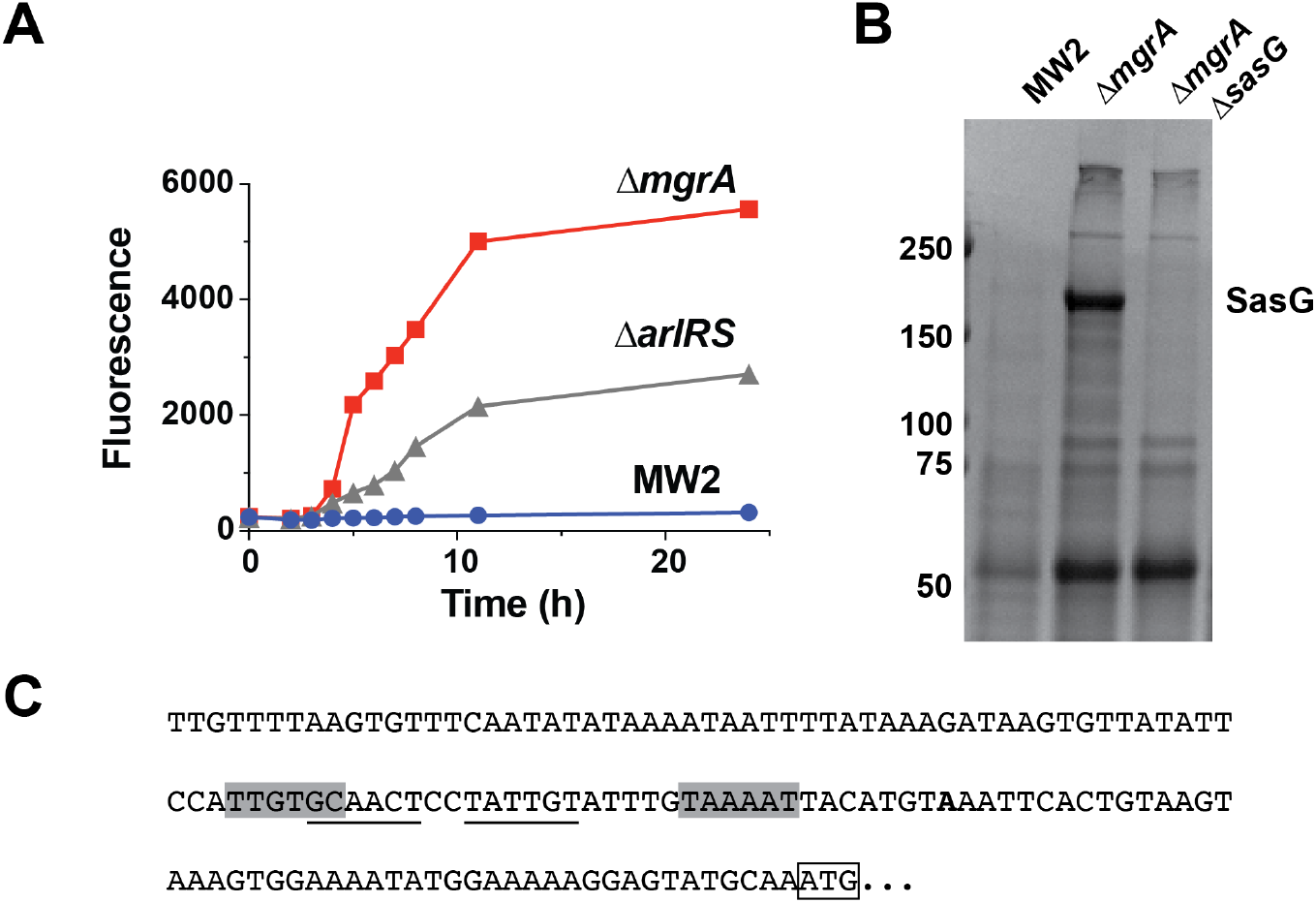
SasG expression is regulated by ArlRS and MgrA. (A) Expression of a P_*sasG*_-GFP transcriptional reporter in the wild-type strain MW2 and isogenic Δ*mgrA* and Δ*arlRS* mutants. (B) Coomassie-stained SDS-PAGE gel of shed surface proteins from MW2, as well as Δ*mgrA* and Δ*mgrA* Δ*sasG* mutants. The SasG band is indicated. (C) Transcription start site (in bold) of *sasG* determined using 5’RACE. The ATG start codon is boxed, and putative -35 and -10 elements are shaded in gray. A potential MgrA binding site is underlined.

### SasG processing after A-domain repeats promotes aggregation in human saliva

As noted in **Figure 1**, a large protein consistent with the size of SasG was upregulated in the Δ*mgrA* mutant and processed to a smaller version after incubation with human saliva (**Figs. 1C & 3A**). These observations suggest that proteases present in saliva could process SasG to smaller sizes. A previous report suggested that SasG possessed self-processing capability and that this cleavage occurred at multiple sites within the B domain [38]. While the self-processing might be occurring in other experimental conditions, we did not observe background processing in our experiments when bacteria were incubated in PBS (**Fig. 1C**). In contrast, our results indicate that SasG may be processed by a host protease(s), and there may be a single cleavage site near one end of the protein, similar to what is seen with Aap [46].

To determine the location of the cleavage site within SasG, we cloned and purified the extracellular portion of SasG. The LPXTG cell wall anchor was replaced with a hexahistidine tag, and the protein was expressed in a *S. aureus* strain that lacks secreted proteases [56]. Purified SasG was incubated with saliva and then re-purified before N-terminal sequencing to determine the cleavage site. The results revealed a cut site after Arg-144, which falls between the A repeats and lectin subdomain (**Fig. 3A**).

**Fig. 3.**
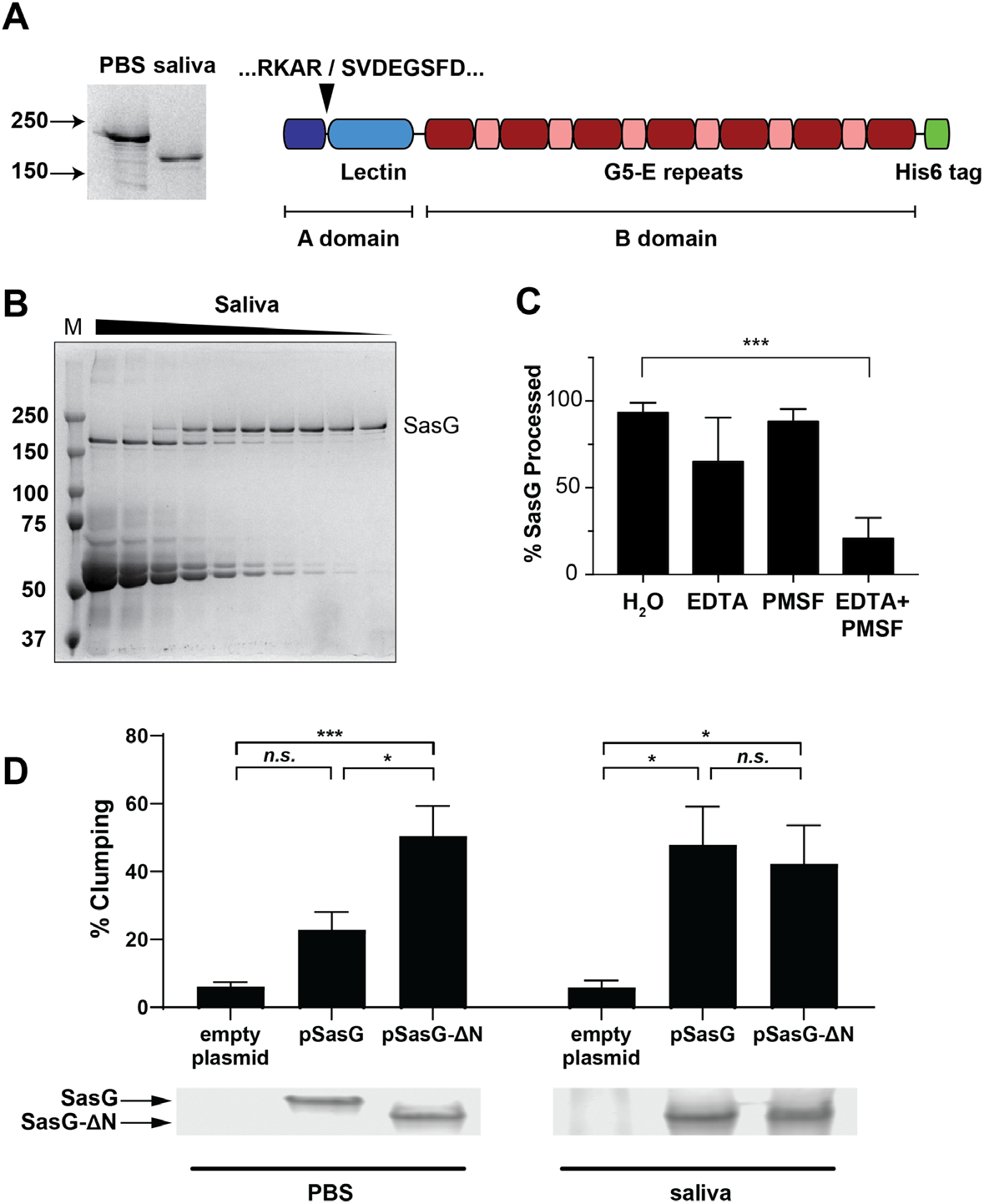
Saliva cleaves SasG within the A domain. (A) The *sasG* gene from *S. aureus* MW2 was cloned with a C-terminal His_6_ tag in place of the cell wall anchor, allowing it to be purified from *S. aureus* culture supernatants. This purified, full-length SasG was then incubated with human saliva for 1.5 h, resulting in SasG cleavage (shown in Coomassie-stained gel on left). Cleaved SasG was re-purified and subjected to N-terminal sequencing, which showed the cleavage site to be N-terminal to the lectin domain. (B) Human saliva was concentrated ∼5-fold before generating a 2-fold dilution series. Purified SasG was then added, and the reactions were incubated for 1 h at 37 °C. (C) Saliva was pre-incubated with either 2.5 mM EDTA, 2.5 mM PMSF, or both, before adding purified SasG. Reactions were incubated for 2 h at 37 °C before resolving on an SDS-PAGE gel. SasG bands were quantified, and the percentage processed to the shorter product was calculated. Results are averages of three experiments, with statistical significance calculated by ANOVA. ***, p < 0.001. (D) Aggregation of LAC strain, lacking its own SasG, and expressing from a plasmid either a full-length SasG construct, or SasG construct with truncated N-terminal domain what replicates the effect of saliva processing. Aggregation was measured on *S. aureus* from overnight cultures suspended in saliva or PBS buffer for 1h. N=7. Coomassie stained SDS-PAGE gels showing expression and processing of SasG constructs in each strain were prepared from cell wall preparations of the above mentioned samples after the incubation.

This is similar to one of the two reported cleavage locations in Aap [46], but it is somewhat surprising because removal of the entire A domain was thought to be required for both Aap and SasG B domain homodimerization and subsequent aggregation [43, 46]. The cleavage of SasG by saliva was found to be dose-dependent (**Fig. 3B**), suggesting presence of specific cleaving protease(s) inside the saliva.

Therefore, purified SasG was incubated with saliva and protease inhibitors to identify the responsible protease(s). Minimal inhibition was seen with EDTA or PMSF alone, but in combination they almost completely inhibited cleavage of SasG (**Fig. 3C**). This result suggests that saliva contains at least two proteases, a metalloprotease and a serine protease, that process SasG and promote bacterial aggregation.

To test if this truncated form of SasG could promote aggregation, we cloned both full-length and truncated versions of *sasG* and expressed them in strain USA300 LAC, which does not express a functional SasG on its own due to a frameshift mutation in its *sasG* gene. While expression of full-length SasG had only minimal effect on aggregation in buffer, and required saliva to facilitate a full-scale aggregation, the truncated version of SasG facilitated aggregation in buffer alone (**Fig. 3D**). This confirmed that removal of the 94 N-terminal amino acids of the A repeat region is sufficient to allow SasG to dimerize and promote aggregation.

### Fractionation to identify host proteases processing SasG

Clarified saliva was concentrated, filtered, and passed over multiple columns to separate the proteins into fractions. First, we used anion exchange chromatography followed by size exclusion chromatography. These fractions were then tested to see if they could cleave purified SasG by running the reactions on SDS-PAGE gels and looking for a shift in SasG size (**Fig. 4A**). The level of SasG cleavage was highest in fractions 19-22 and these fractions were used going forward. In parallel, we tested the response of the isolated active saliva fraction with protease inhibitors to determine the exact class of the enzyme. The most inhibition was observed with AEBSF, Antipain, and Leupeptin, suggesting the enzyme present in the active fractions is a serine protease (**Fig. 4B**). After electrophoresis separation of fractions with highest activity (**Fig. 4A**), individual bands were extracted from the gel and the protein(s) identified by MALDI mass spectrometry. Seven proteases were detected in these bands with significant peptide coverage, including trypsin-1, prostasin, serine protease 27 and various cathepsins (**Supp. Table 1**). Considering the protease inhibitor patterns (**Fig. 4B**), the best hit from the proteomics assessment was human trypsin.

**Fig. 4.**
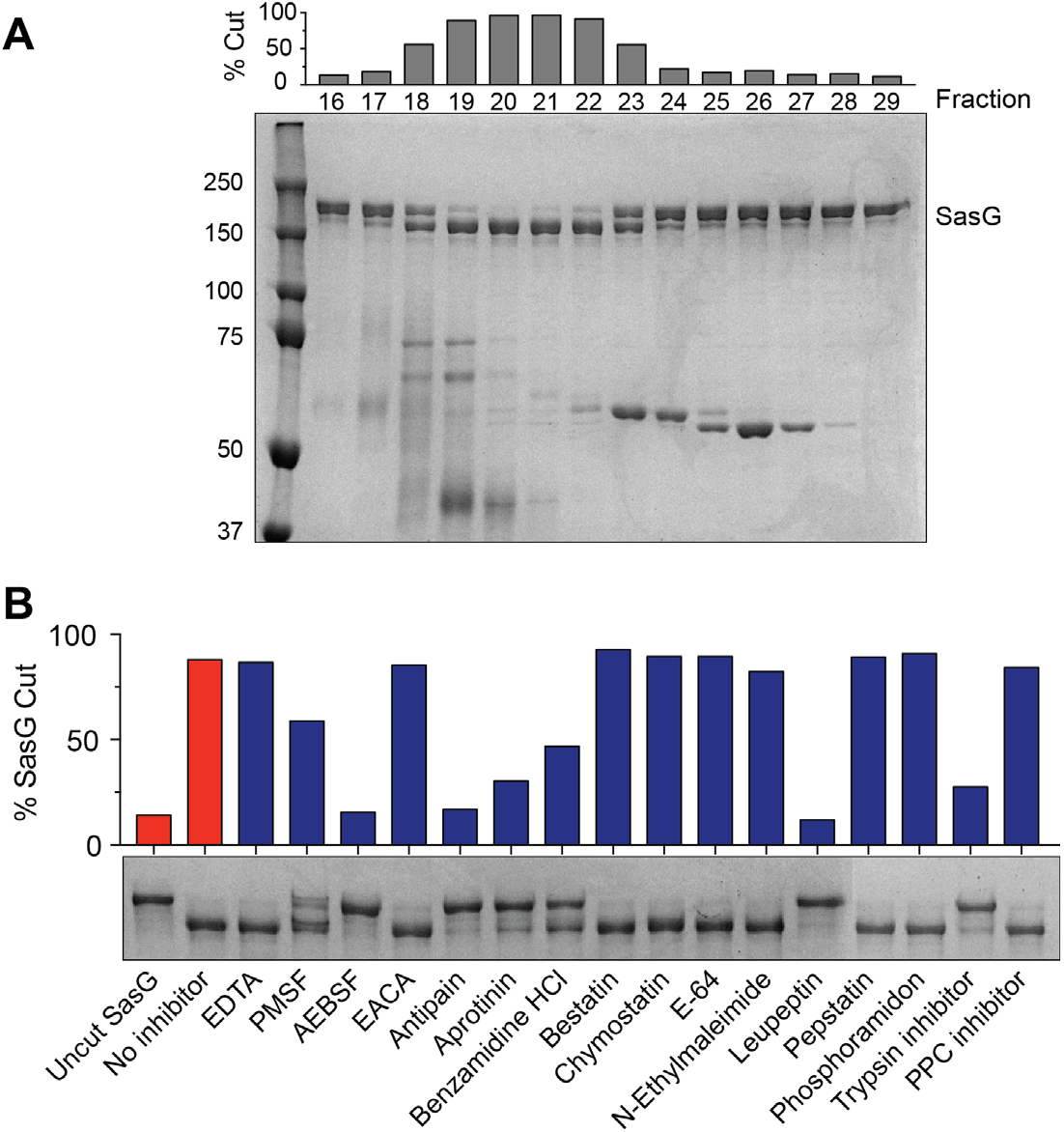
Partial purification of SasG processing enzyme from human saliva. (A) Pooled active fractions from passing saliva over an anion exchange column were then passed over a size exclusion column. Coomassie stained gel shows SasG cleavage by selected fractions from the size exclusion purification. Fraction numbers are indicated above the gel, and bars show percent SasG cleavage for each fraction. Molecular weight standards in kDa are indicated on the left. (B) Aliquots of fraction 20 were pre-incubated with the indicated protease inhibitors for 15 min before adding SasG. Cleavage of SasG was measured after 1.5 h at 37°C by separating on an SDS-PAGE gel and quantifying percent cleavage.

### Validation of identified proteases

We used commercially available trypsin to test SasG processing and promotion of *S. aureus* aggregation. A range of trypsin concentrations (0-200 μg/mL) was incubated with purified SasG, and dose-dependent SasG processing was visualized on SDS-PAGE (**Fig. 5A**). In parallel we performed aggregation assays at the same doses of protease (**Fig. 5B**). At 0.2 μg/mL we started observing cleavage of SasG, which correlated with an increase in aggregation. The levels of cleavage and aggregation increased at 2 μg/mL trypsin and remained fairly constant at 20 μg/mL (**Fig. 5A, B**). These findings demonstrated that trypsin can recapitulate the phenotype of SasG processing and promote aggregation. At 200 μg/mL trypsin, the whole SasG protein was becoming degraded (**Fig. 5A**), and the aggregation phenotype was mostly lost (**Fig. 5B**).

**Fig. 5.**
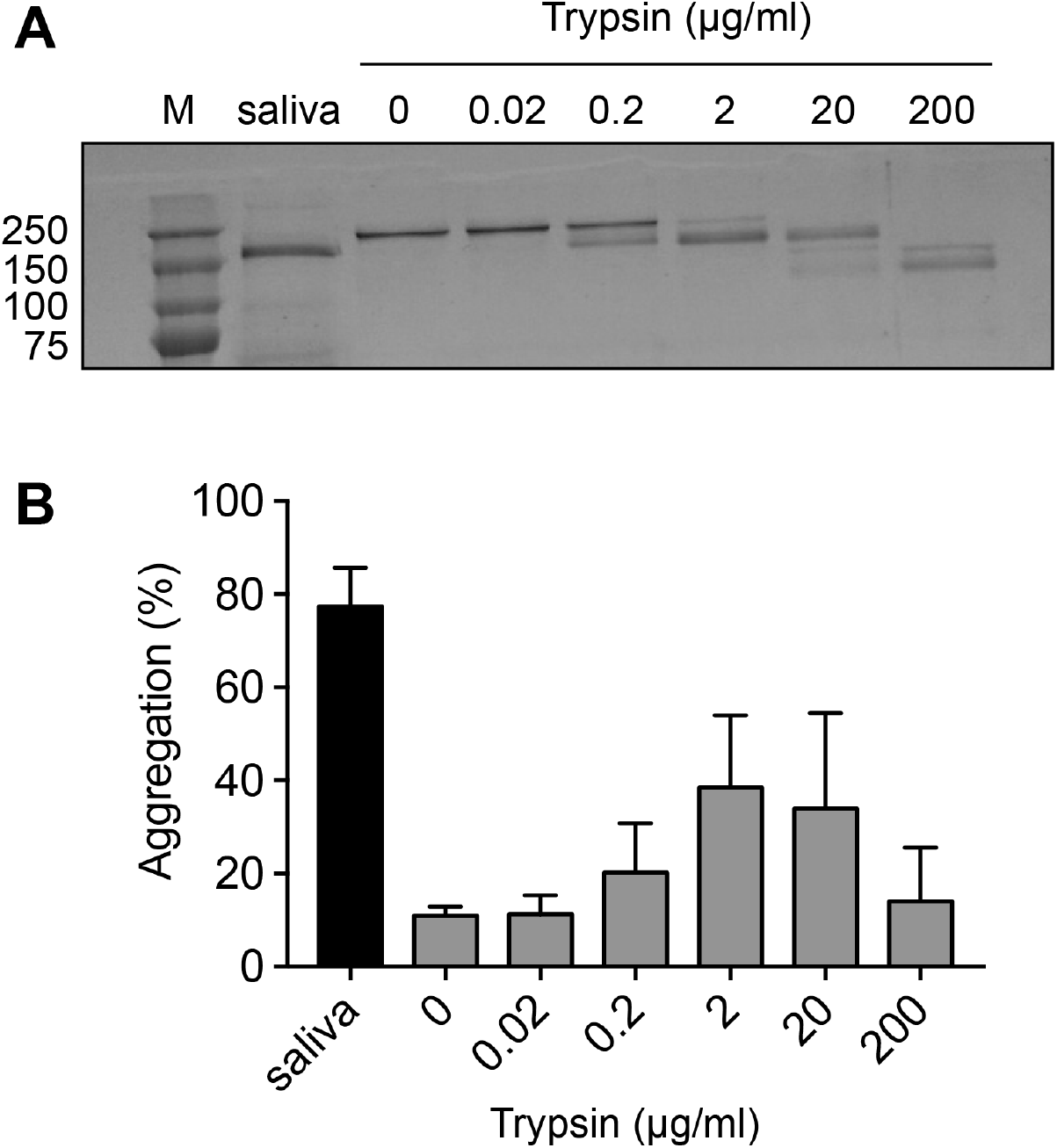
Trypsin can process SasG and promote *S. aureus* aggregation. (A) Purified full-length SasG was incubated for 1 h with either human saliva or serial dilutions of trypsin before running on an SDS-PAGE gel and staining with Coomassie. (B) *S. aureus* MW2 Δ*mgrA* cells were resuspended in either saliva or PBS supplemented with trypsin and allowed to aggregate for 1 h. Measurements are averages and standard deviations of three separate experiments.

### Role of SasG in pneumonia model

To examine the biological relevance of SasG *in vivo*, we intratracheally infected mice with MW2 Δ*mgrA* (thus, SasG-expressing) or with Δ*mgrA* Δ*sasG* double mutant (**Fig. 6**). No evidence of systemic dissemination was observed in this model (**Fig. 6A**). The mice that were infected with the double mutant lacking SasG showed decreased number of colonies in the lungs (**Fig. 6B**), compared to the Δ*mgrA* strain expressing SasG. At the same time markers of inflammation and tissue damage, that is number of leukocytes (**Fig. 6C**) and level of protein (**Fig. 6D**) in the bronchoalveolar lavage (BAL) remain similar irrespective of the injected strain. The same trend of decreased bacterial counts and not significantly affected leukocytes and protein levels was also observed when a lower dose of *S. aureus* was used for infection (**Supplementary Fig. 1A-C**).

**Fig. 6.**
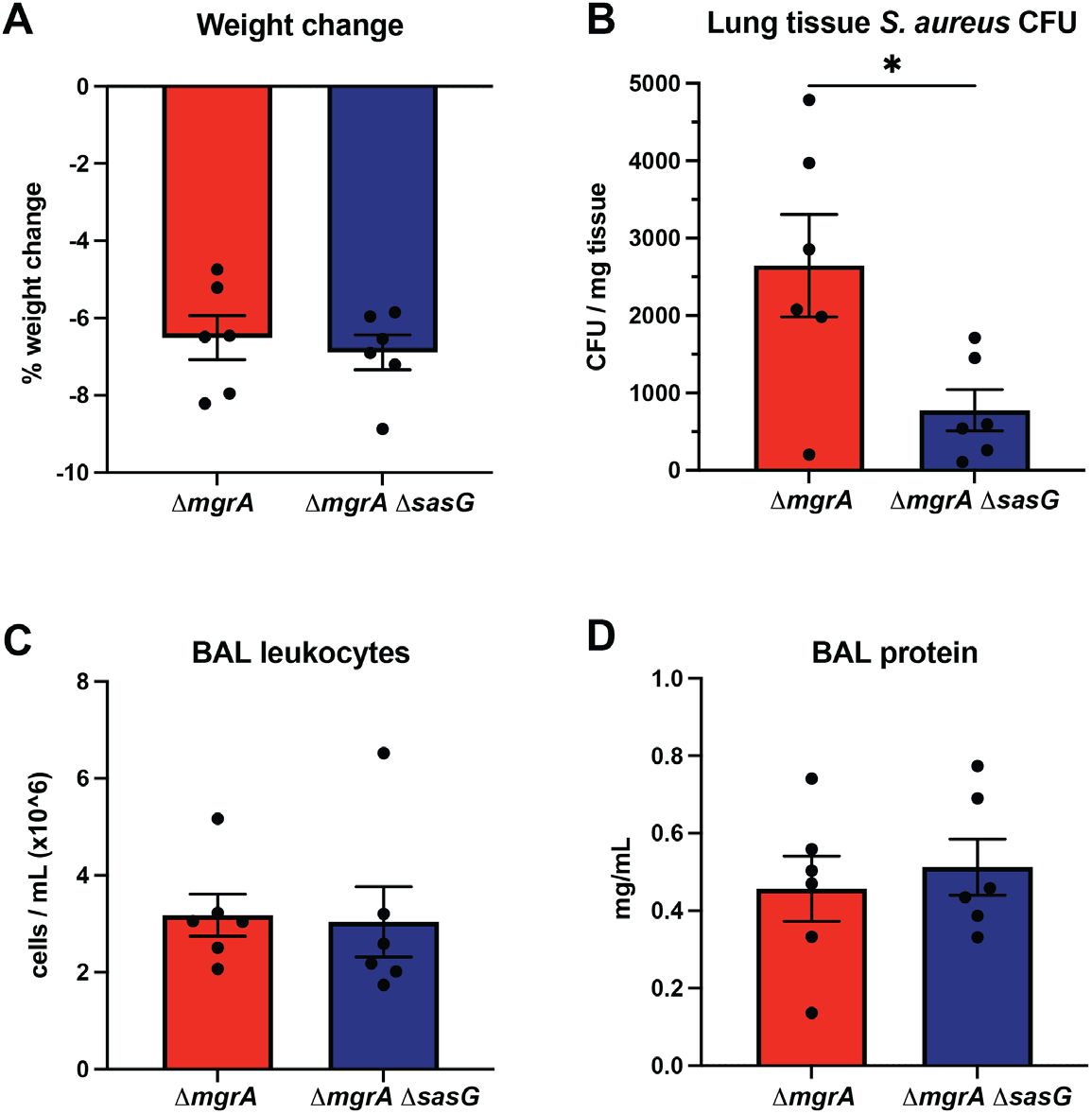
SasG is involved in *S. aureus* virulence in lung infection. Mice were infected intratracheally by *S. aureus* MW2 Δ*mgrA* and by its congenic strain Δ*mgrA* Δ*sasG* lacking SasG, and severity of pneumonia was assessed by weight loss (A), counting the CFU burden in lung homogenates (B), lung leukocyte recruitment in bronchoalveolar lavage (C), and protein infiltration in lavage fluid (D) after 24h. Results presented as means ± SEM, with statistical significance calculated by Mann-Whitney test. *, p < 0.05.

**Fig. 7.**
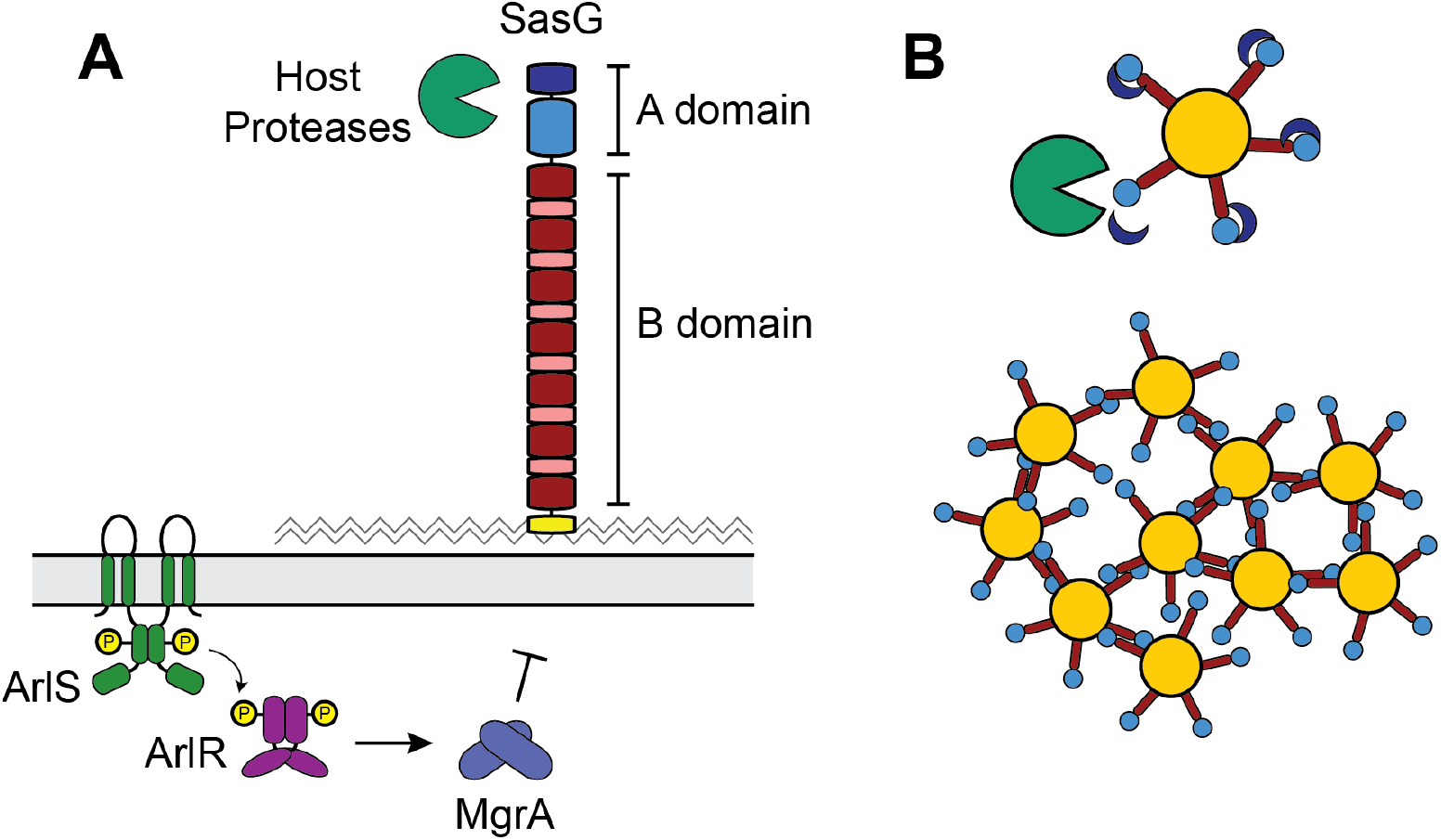
Model of SasG transcriptional and post-translational regulation. (A) Expression of *sasG* is repressed by the ArlRS-MgrA regulatory cascade. At the post-translational level, host proteases such as trypsin can remove the N-terminal end of the A domain. (B) Removal of the end of the A domain (dark blue) allows SasG to oligomerize with SasG molecules on neighboring cells, resulting in aggregation of *S. aureus*.

Overall, this suggests that during lung infection the presence of SasG on *S. aureus* surface has no effect on host response or local damage, but it does benefit survival of the pathogen when faced with host immune response. Overall, the mouse pneumonia data indicate that presence of SasG contributes to *S. aureus* virulence *in vivo*.

## Discussion

Roughly one-third to half of healthy individuals are colonized by *S. aureus* in the nasal cavity and/or nasopharynx [57-59]. While *S. aureus* colonization is benign in healthy adults, presence of *S. aureus* in the respiratory tract is the major risk factor for developing pneumonia in the intensive care unit [60, 61]. Despite the high rate of *S. aureus* carriage in the oral cavity, only preliminary studies have been performed with *S. aureus* interactions with human saliva proteins [62, 63]. *S. aureus* predominantly binds human proteins using microbial surface components recognizing adhesive matrix molecules (MSCRAMMs) [64]. We have previously shown that the ArlRS/MgrA regulatory cascade controls expression of MSCRAMMs and other surface proteins that function in adhesion and immune evasion [65]. Strains lacking either *arlRS* or *mgrA* overexpress these surface proteins, and in this work we made the surprising discovery that a *S. aureus mgrA* mutant aggregates in the presence of human saliva. We found that intercellular aggregation is dependent on expression of SasG, but also requires host factors in saliva to process SasG.

In previous studies we demonstrated that full-length SasG is sufficient to block clumping and adhesion of cells by physically interfering with other surface proteins’ ability to bind to host matrix components [49, 51, 66, 67]. However, SasG expression is low in *S. aureus* laboratory strains under standard *in vitro* conditions, which masks these clumping interference and aggregation phenotypes. Through our mapping of the *sasG* promoter and transcriptional reporter assay, we show that *sasG* expression is repressed by ArlRS/MgrA, and we identify a potential MgrA binding site that overlaps with the *sasG* promoter. Therefore, inactivation of the ArlRS-MgrA cascade allows for high expression levels of *sasG*.

We also found that there is significant variation in *sasG* expression and molecular characteristics among strains: not all *S. aureus* strains have a functional (full-length) copy of *SasG*, and of the strains that have the functional gene, not all express SasG at detectable levels. USA400 MW2 and 502a encode full length, surface-attached copies of SasG with 5 B-repeats which aggregate with high efficiency. Bioinformatic analysis of the CF isolate AH4654 genome revealed the *sasG, mgrA*, and *arlRS* genes, and their respective promoter regions are all essentially identical to MW2. Interestingly, this CF isolate expresses SasG and aggregates natively (**Fig. 1**), similar to other *S. aureus* isolates that that fall into ST15/CC15 grouping [47]. In contrast strain Newman, despite encoding a full-length SasG, does not present it on its surface and does not aggregate with or without MgrA. The reason SasG is not functional in Newman is unclear at this time. Strains such as USA300 LAC and N315 have truncated copies of SasG due to frameshift mutations and therefore cannot aggregate. Other strains like MN8 and MRSA252 do not possess *sasG* and any observed aggregation was likely due to another surface protein.

SasG is one of the key drivers of biofilm formation in *S. aureus* [38, 40, 45, 47]. SasG, and its *S. epidermidis* homolog Aap, consist of multiple domains with distinct functions (**Fig. 1A**). In *S. epidermidis* the Aap A domain is known to be removed by the secreted metalloprotease SepA to facilitate biofilm accumulation [46], but native *S. aureus* secreted proteases have not been found to cleave SasG in the same manner [38]. Previous studies on *S. epidermidis* Aap also showed that exogenously added host proteases, such as trypsin and cathepsin G, could cleave Aap and enhance biofilm formation through processing [43]. Our studies have found a parallel role for host proteases in cleaving *S. aureus* SasG and triggering aggregation.

During infection, *S. aureus* uses mechanisms of aggregation and biofilm formation as survival strategy to protect itself long-term in response to environmental stressors, such as antimicrobials or host immune factors. Our data demonstrates that upregulation of *sasG* is associated with increased aggregation upon interaction with human saliva, which is known to contain numerous proteases [68]. Considering that the aspiration of saliva secretions is a common precursor to lung infection [69], our findings indicate that salivary proteases are capable of cleaving SasG at a single site within the A domain. This processing removed the 94 amino acids that compose the A-repeats, exposing the A-lectin and B-domains to interact on neighboring cells and homodimerize. We fractionated the proteases to identify human trypsin and validated with commercially available trypsin. However, additional serine and metalloproteases may also contribute to processing of SasG. From an adaptive standpoint, *S. aureus* may have evolved a surface protein like SasG that is proteolytically labile, which can sense environmental conditions and facilitate aggregation to protect *S. aureus* under stress.

Despite significant biochemical and structural studies on SasG, accompanied by experiments *in vitro*, there are no studies determining its contribution to virulence in animal models of infection. However simultaneous deletion of SasG and Eap did reduce insect mortality in a silkworm infection model [70]. In this work, we provide evidence that SasG contributes to *S. aureus* in establishment of a lung infection. We demonstrated that SasG is important for *S. aureus* to survive and proliferate at the infection site.

However, the presence of SasG did not impact the host response or damage to host, suggesting it is solely important for *S. aureus* survival in a stressful environment.

In summary, we have shown that the global regulator MgrA controls expression of the surface protein SasG. There is variation in the type and amount of *sasG* expressed among *S. aureus* strains, but expression of full-length SasG is associated with increased aggregation which is dependent on the presence of host proteases. We identified the serine protease human trypsin as a component of saliva that can process SasG A-domain to trigger aggregation. Finally we showed that SasG is important for full virulence in a *S. aureus* lung infection.

## Material and Methods

### Reagents and growth conditions

*S. aureus* strains and plasmids used in this work are listed in **Table 1**. AH4654 is one of 75 clinical isolates, isolated from 10 pediatric CF patients and kindly gifted by the Starner Lab, University of Iowa. *S. aureus* was cultured in tryptic soy broth (TSB) or brain heart infusion (BHI) broth, and *E. coli* was cultured in lysogeny broth (LB) at 37 °C with shaking at 200 rpm. Antibiotics were added to the media at the following concentrations: chloramphenicol (Cam), 10 μg/mL; erythromycin (Erm), 5 μg/mL; and tetracycline (Tet), 1 μg/mL. *E. coli* strains with plasmids were maintained on media supplemented with ampicillin at 100 μg/mL; kanamycin, 50 μg/mL; or spectinomycin at 50 μg/mL. Porcine trypsin and the Protease Inhibitors Set (Roche) were purchased from Sigma. Stimulated saliva was collected over 10-30 min by chewing on paraffin wax. Particulate material was removed by centrifugation, and this clarified saliva was stored at 4°C for up to 2 days.

**Table 1.** Bacterial strains and plasmids

| Strain/plasmid | Genotype/properties | Reference |
| --- | --- | --- |
| *E. coli* |  |  |
| DH5α | Cloning strain | Protein Express |
| DC10B | Cloning strain (*dcm*<sup>-</sup>) | [71] |
| T7 Express | Protein expression strain | NEB |
| *S. aureus* |  |  |
| RN4220 | Restriction deficient cloning host | [80] |
| MW2 | USA400 CA-MRSA | [81] |
| AH3422 | MW2 $\Delta mgrA$ | [49] |
| AH3989 | MW2 $\Delta mgrA \Delta sasG$ | [49] |
| AH1263 | USA300 CA-MRSA Erm<sup>S</sup> (LAC<sup>*</sup>) | [82] |
| AH3375 | LAC $\Delta mgrA$ | [49] |
| AH1919 | LAC<sup>*</sup> $\Delta aur \Delta sspAB \Delta staphopainA \Delta spl::erm$ | [56] |
| AH4607 | LAC<sup>*</sup> $\Delta aur \Delta sspAB \Delta staphopainA \Delta spl::erm \phi 11 att::tet$ | This work |
| 502a | ST5 MSSA |  |
| AH3625 | 502a $\Delta mgrA::tetM$ | [49] |
| Newman | MSSA | [83] |
| AH3472 | Newman $\Delta mgrA::tetM$ | [49] |
| N315 | USA100 MRSA | [84] |
| AH3473 | N315 $\Delta mgrA::tetM$ | [49] |
| MN8 | USA200 MSSA | [85] |
| AH3480 | MN8 $\Delta mgrA::tetM$ | [49] |
| MRSA252 | USA200 HA-MRSA | [86] |
| AH3483 | MRSA252 $\Delta mgrA::tetM$ | [49] |
| AH4654 | MSSA CF Isolate | This work |
| AH4728 | AH4654 $\Delta sasG::Tn Erm$ | This work |
| **Plasmids** |  |  |
| pALC2073 | Tetracycline-inducible shuttle vector, Cam<sup>R</sup> | [87] |
| pRMC2 | Tetracycline-inducible shuttle vector, Cam<sup>R</sup> | [88] |
| pCM28 | Empty vector control for pCM29, Cam<sup>R</sup> | [82] |
| pCM29 | sGFP expression vector, Cam<sup>R</sup> | [78] |
| pTEV5 | Expression vector with TEV-cleavable His6 tag, Amp<sup>R</sup> | [89] |
| pHC66 | *mgrA* complementation vector, Cam<sup>R</sup> | [49] |
| pHC89 | pALC2073-*sasG* | This work |
| pHC90 | pALC2073-*sasG*-His6 (secreted) | This work |
| pHC108 | pTEV5 *sasG* B repeat | This work |
| pHC116 | pALC2073-*sasG* $\Delta N$ | This work |
| pHC127 | P<sub>*sasG*</sub>-sGFP, Cam<sup>R</sup> | This work |

### Recombinant DNA and genetic techniques

*E. coli* DH5α and DC10B were used as a cloning host for plasmid constructions. Restriction enzymes, DNA ligase, and Phusion DNA polymerase were purchased from New England Biolabs. The plasmid mini-prep and gel extraction kits were purchased from Invitrogen. *S. aureus* genomic DNA was purified using the Puregene yeast/bacteria kit B (Qiagen). Lysostaphin, used for *S. aureus* DNA extractions, was purchased from Sigma. Plasmids were purified from *S. aureus* RN4220 or *E. coli* DC10B and electroporated into MRSA LAC strains as described previously [71, 72].

Bacteriophage transductions between *S. aureus* strains were performed with phage 11 as described previously [73]. All oligonucleotides were ordered from IDT (Coralville, IA) and are listed in **Table 2**. Routine DNA sequencing was performed at the University of Iowa DNA Core Facility or the Molecular Biology Service Center at the University of Colorado Anschutz Medical Campus. Whole genome sequencing was performed at the University of Iowa DNA Core Facility with the Illumina MiSeq platform followed by *de-novo* contig generation with the SPAdes genome assembler [74], and quality assessed with QUAST [75]. Assemblies were annotated with Prokka [76]w. The draft genome of AH4654 was deposited to NCBI and Illumina data is available in Genbank (accession no. JAPQKW000000000).

**Table 2.**
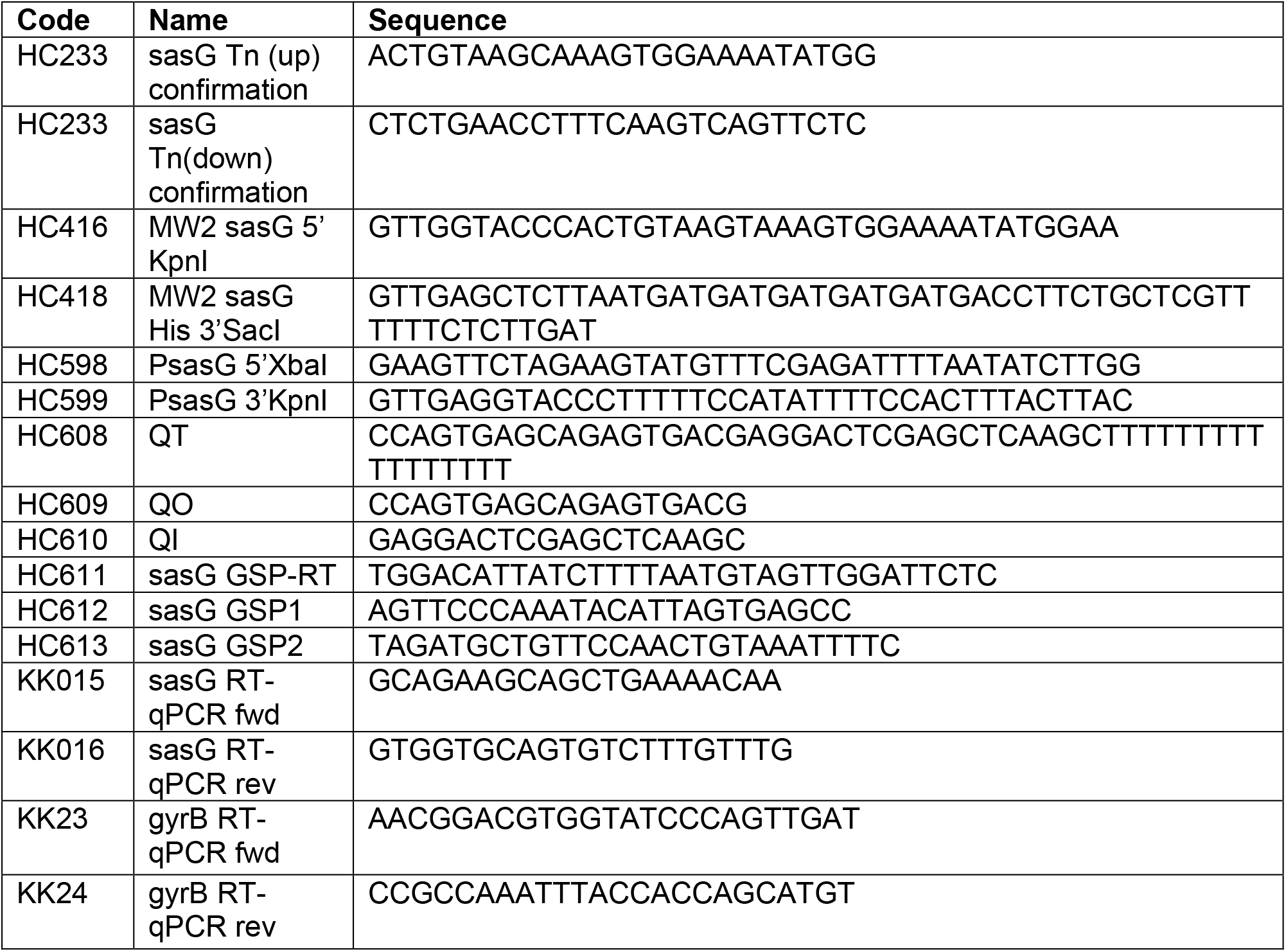
Primers

### RNA purification and RT-qPCR

Bacterial cultures were grown overnight in TSB and then subcultured to an OD_600_ of 1.5. Cells were then pelleted and washed with RNAprotect Bacterial Reagent (Qiagen). To extract RNA, cells were lysed with lysostaphin for 30 minutes at room temperature, and RNA was purified using the RNeasy Mini Kit (Qiagen). Following RNA purification, genomic DNA was then removed using the Turbo DNase Kit (Ambion). cDNA was then generated from DNase treated RNA template using the iScript cDNA synthesis kit (Bio-Rad). To perform quantitative PCR (qPCR), Primers KK15 and KK16 were used for *sasG*, and KK23 and KK24 for DNA gyrase *(gyrB)*, as described previously [49]. qPCR was performed by amplifying cDNA in 20 μL reaction volumes with iTaq Universal SYBR Green Supermix (Bio-Rad) in the CFX96 Touch Real-Time PCR System (Bio-Rad) under the following conditions: 3 min at 95°C, 40 cycles of 10 s at 95°C and 30 s at 59°C, followed by a dissociation curve. No template and no reverse transcription controls were performed in parallel. Experiments were performed in biological triplicate with two technical replicates, expression was normalized to *gyrB*.

### *sasG* promoter mapping and GFP fusion plasmid

The *sasG* promoter was mapped using rapid amplification of 5’ cDNA ends (5’ RACE) [77]. Template RNA was purified from MW2 Δ*mgrA* using the RNeasy Mini Kit (Qiagen) as previously described [49]. Primers used were the general 5’ RACE primers [77] HC608, HC609, and HC610, and the *sasG*-specific primers HC611, HC612, and HC613. To generate the P_*sasG*_-GFP fusion plasmid, the region upstream of *sasG* was amplified using primers HC598 and HC599. The fragment was digested using XbaI and KpnI before ligating into pCM29 [78]. The resulting plasmid, pHC127, encodes the *sasG* promoter upstream of an optimized ribosome binding site and codon optimized gene for superfolder GFP. To assess expression, overnight cultures were diluted 1:100 in TSB containing chloramphenicol in a black 96-well plate. Plates were incubated at 37 °C with shaking in a humidified microtiter plate shaker (Stuart). A Tecan Infinite M200 plate reader was used to periodically measure OD_600_ and fluorescence intensity with excitation at 495 nm and emission at 515 nm. Values represent averages and standard deviations of triplicate wells.

### *S. aureus* aggregation assay

*S. aureus* cultures (5 mL) were grown overnight in TSB with shaking at 37 °C. One mL of culture was harvested by centrifugation and the media was discarded. The cells were resuspended in 1 ml of either phosphate buffered saline or clarified human saliva. Tubes were allowed to sit for 1 h at room temperature, and then aggregation was visually assessed. For quantification of aggregation, 100 μL of liquid was removed from the top of the tube at 0 h and 1 h, and the optical density at 600 nm was measured in a 96-well plate in a Tecan infinite M200 plate reader. Measurements represent averages and standard deviations of experiments performed on three separate days.

### Cell wall preparations

For preparation of cell wall proteins after aggregation assays, the tubes were centrifuged, and the cells were washed twice with PBS. The cells were resuspended in 500 μL of protoplasting buffer (10 mM Tris pH 8, 10 mM MgSO_4_, 30% raffinose).

Lysostaphin (25 μg) was added and the cells were incubated for 1 h at 37 °C. The tubes were centrifuged for 3 min at max speed, and 500 μL of supernatant was transferred to a new tube. Proteins were precipitated by adding 125 μL of cold trichloroacetic acid and leaving on ice for 2 h. Precipitated proteins were pelleted by centrifuging at max speed for 10 min. The pellet was washed twice with 500 μL of cold 100% ethanol and then inverted to dry. The pellets were resuspended in 36 μL of SDS-PAGE loading dye, heated to 85°C, and then 10 μL was loaded on a 7.5% acrylamide gel.

### Purification of full-length SasG

The *sasG* gene from *S. aureus* MW2 was amplified using primers HC416 and HC418 (**Table 2**), which remove the last 33 amino acids of SasG, including the LPXTG cell wall anchor, and replace them with a glycine followed by six histidine residues. This C-terminally tagged, secreted version of *sasG* was cloned into pALC2073 under the control of an anhydrotetracycline-inducible promoter, generating pHC90. We decided to purify this version of SasG from *S. aureus* LAC, which does not have an intact copy of *sasG* on the chromosome. To avoid potential proteolysis, we used a previously developed strain of LAC lacking secreted proteases (AH1919). Additionally, we modified AH1919 to be resistant to anhydrotetracycline by integrating the empty vector pLL29 [79] in the phage 11 attachment site, generating host strain AH4607.

For expression of SasG, pHC90 was moved into AH4607 and a 5 mL culture was grown overnight at 37 °C in TSB with chloramphenicol. This overnight culture was used to inoculate 1 L of TSB supplemented with chloramphenicol and 0.15 μg/mL anhydrotetracycline. The culture was grown with shaking for ∼6.5 h at 37°C. Cells were removed by centrifugation, and the culture supernatant was concentrated to ∼30 mL using an Amicon stirring pressure concentrator with a 100 kDa cutoff filter. The supernatant was dialyzed twice against binding buffer (50 mM sodium phosphate, 300 mM NaCl, pH 8). SasG-His6 was then purified using a pre-packed 5 ml IMAC cartridge (Bio-rad) on a Bio-rad FPLC. SasG-His6 was eluted with a linear gradient up to 100% elute buffer (50 mM sodium phosphate, 300 mM NaCl, 250 mM imidazole, pH 8). The protein was then concentrated and dialyzed against storage buffer (20 mM sodium phosphate, 150 mM NaCl, pH 7.5). Glycerol was added to 20% before flash freezing and storing at -80°C.

### SasG processing assays

Purified, full-length SasG was diluted 10-fold in phosphate buffered saline, and 2 μL of this dilution was combined with 2 μL of water and16 μL of clarified saliva or saliva fraction. Reactions were incubated for 1 h at 37 °C unless otherwise indicated.

Processing was then quenched by adding 7 μL of SDS-PAGE loading buffer and heating to 65 °C. 10 μL of this was then loaded on a 7.5% or 10% gel, or a 4-20% gradient gel. For calculating the percentage of SasG processed, Coomassie-stained gels were scanned and quantified using Image Studio Lite (LiCor).

### Identifying the cleavage site within SasG

A large SasG cleavage reaction was setup using 100 μL of purified SasG-His6, 900 μL of PBS, and 4 ml of clarified, filtered saliva. The reaction was allowed to incubate for 1.5 h at 37 °C. The solution was then exchanged to binding buffer (same as above) using a 100 kDa molecular weight cutoff filter (Amicon). SasG-His was then re-purified using HIS-Select resin (Sigma) and eluted with bind buffer containing increasing concentrations of imidazole. Fractions containing SasG-His were pooled and concentrated to ∼0.5 mL, and 2, 4, 6, and 8 μL aliquots were mixed with SDS-PAGE buffer, boiled, and run on a 4-15% gradient gel. Proteins were then transferred to a PVDF membrane using a Trans-blot Turbo transfer system (Bio rad) and the membrane was stained with Coomassie. N-terminal sequencing of cleaved SasG was carried out by Edman degradation using a Shimadzu PPSQ-53A Gradient Protein Sequencer at the Protein Facility at Iowa State University.

### Partial purification of proteases from human saliva

Stimulated saliva (∼90 ml) was collected over one day and centrifuged at 30,000* g to remove debris. Clarified saliva was filtered and then concentrated to ∼3 mL using 30,000 MWCO centricon concentrators (Amicon) and dialyzed against buffer A (20 mM Tris pH 8, 2 mM NaCl). The sample was then separated by anion exchange chromatography using a HiScreen Capto Q column (GE Life Sciences), eluting with a linear gradient up to 100% buffer B (20 mM Tris pH 8, 1 M NaCl). Fractions were tested using the SasG processing assay described above, except that tubes were incubated for 2 h at 37 °C before running on an SDS-PAGE gel. Active fractions were pooled, concentrated to ∼350 μL, and loaded on an SEC70 size exclusion column (Bio-rad).

The running buffer consisted of 20 mM Tris pH 8 and 100 mM NaCl. 0.5 ml fractions were collected and tested for their ability to cleave SasG as described above, and a couple fractions (20 and 21) was selected for further analysis. Protease inhibitors (Sigma) were used according to manufacturer’s instructions. For protein identification, bands were excised from an SDS-PAGE gel and analyzed at the University of Colorado Mass Spectrometry Proteomics Shared Resource Facility.

### Pneumonia model

All mouse experiments were conducted in accordance with National Institutes of Health guidelines and previously approved by the University of Colorado Institutional Animal Care and Use Committee. Wild-type (WT) female BALB/c, 6-8 weeks old, were purchased from Jackson Laboratories (Bar Harbor, ME). Mice were anesthetized with isoflurane inhalation and challenged with approximately 2*10^8^ colony-forming units (CFU)/30 μL of either mutant (Δ*mgrA* or Δ*mgrA* Δ*sasG*) *S. aureus* MW2 strain intratracheally. A blunt-tipped, bent 18-g Hamilton syringe was used to administer 30 μL of *S. aureus* directly into the lungs. Mice were left to recover for 24 hours after which were euthanized using lethal dose of ketamine/xylazine. Trachea was cannulated and the right lobes were tied off allowing for unilateral bronchial alveolar lavage (BAL) fluid isolation from the left lung. The right lobes were weighed then homogenized for CFU determination. As a measure of lung inflammation and injury, leukocytes and protein in BAL fluid were measured.

## Acknowledgments

We thank Dr. Timothy Starner for providing CF isolate AH4654. H. Crosby was supported by American Heart Association postdoctoral fellowship 15POST25720016. K. Keim was supported by the NIAID Molecular Pathogenesis of Infectious Disease T32 predoctoral fellowship AI052066-19. Research in the laboratory of A. R. Horswill was supported by NIH grants AI083211 and AI162964.

